# Single-cell transcriptomics of enriched rare neural lineages of *Tribolium castaneum*

**DOI:** 10.64898/2026.11.03.763907

**Authors:** Noel Cabañas, Ana Veloso, Robert Zinzen, Gregor Bucher, Christine Mau

## Abstract

**Background:** Single-cell transcriptomic atlases have transformed our understanding of cell diversity. However, studies designed to profile entire tissues or organisms often omit or underrepresent rare cell populations. Enriching selected populations before sequencing offers a powerful approach to overcoming this limitation, but its application in insects has largely focused on *Drosophila*, limiting opportunities for comparative studies. Here, we establish complementary workflows for targeted transcriptomic profiling of rare neural populations in the red flour beetle, *Tribolium castaneum*.

**Results:** We developed two workflows combining population enrichment with single-cell or single-nucleus RNA sequencing. First, we generated cell suspensions from a transgenic reporter line labelling embryonic neuroblasts, enriched fluorescently labelled cells by fluorescence-activated cell sorting, and performed single-cell RNA sequencing. Second, we immunolabelled first-instar larval brains with an antibody against the nuclear transcription factor Rx, isolated and sorted labelled nuclei, and performed single-nucleus RNA sequencing. Both workflows yielded material suitable for transcriptomic profiling, demonstrating complementary reporter-based and antibody-based strategies for accessing selected neural populations at different developmental stages.

**Conclusions:** These workflows establish a methodological framework for targeted single-cell and single-nucleus transcriptomic studies of rare or specific neural populations in *Tribolium*. They provide practical starting points for adaptation to other insect systems, expanding opportunities for comparative investigations of neural development and cell-type evolution.

## Background

Single-cell omics methods and technologies have changed how we see cell diversity and its molecular landscape. We are now able to comprehensively profile small complete organisms, organs, tissues, or organoids to uncover their cell types and dynamic heterogeneity (Baysoy et al., 2023; Li, 2021; Liu and Li, 2024; Sun et al., 2023; Vandereyken et al., 2023; Wang et al., 2024; Wen et al., 2022; Wu et al., 2024). For example, single-cell transcriptomics has been used to explore gene expression in a variety of organs and organisms, including many insects in the orders Diptera, Coleoptera, Lepidoptera, Hymenoptera, and Hemiptera (Arya et al., 2024; Kwon et al., 2021; Li, 2021; Liu and Li, 2024; Liu et al., 2024; Loh et al., 2025; Page et al., 2023; Prakash et al., 2024; Robben et al., 2024; Rossi et al., 2026; Sun et al., 2023; Wan et al., 2026) and some insect outgroups in the orders Araneae, Decapoda, and Cladocera (Cui et al., 2024; Jin et al., 2023; Koiwai et al., 2021; Krishnan et al., 2024; Leite et al., 2024; Liu and Li, 2024; Medina-Jiménez et al., 2024; Xin and Zhang, 2023; Yang et al., 2022; Zhong et al., 2026). These reference single-cell atlases enable rapid, effective identification of cell types and their expression profiles for comparative single-cell transcriptomic studies, driving significant advances in defining cell types, states, trajectories, spatial contexts, and plastic responses (Kamimoto et al., 2023; MacLean et al., 2018; Saunders et al., 2023; Vandereyken et al., 2023). However, these resources are biased towards the most abundant cells and underrepresent or omit rare cell types, making it difficult to address evolutionary and developmental questions in specific cell types in detail. An alternative strategy based on the enrichment of labelled cells is more suitable and advantageous for the precise isolation of specific neural cell types to enhance the resolution of single-cell datasets. Fluorescence-activated cell sorting (FACS) of marked cells has been extensively reported (Fabian et al., 2022; Hammond et al., 2019; Kurmangaliyev et al., 2019; McLaughlin et al., 2021; McLaughlin et al., 2022a; Ratz et al., 2022; Severo et al., 2018; Vöcking and Famulski, 2023), but sorting coupled with single-cell sequencing has not been widely adopted in any other insect other than *Drosophila*. In flies, FACS enrichment followed by single-cell sequencing has been reported mainly in larval and pupal stages. While in embryos, only a handful of examples exist for heart progenitor cells, motoneurons, salivary glands, muscle cells, and one more on NBs (Cabañas et al., 2026; Huang et al., 2023; May and Röper, 2024; Secchia et al., 2022; Velten et al., 2022; Vogler et al., 2021). In other insects, this strategy has been reported only in two insects, *Anopheles gambiae* and *Tribolium castaneum* (Cabañas et al., 2026; Page et al., 2023; Severo et al., 2018). This is a clear knowledge gap for developmental comparative studies in rare insect cells.

All aforementioned studies relied on antibodies or transgenic lines to label the cells of interest prior to sorting. However, these resources are time-consuming to establish and are not always available for non-model organisms, limiting the capacity to perform comparative studies across insect cell lineages. To address this issue, we present the red flour beetle, *Tribolium castaneum,* as a model for neural comparative single-cell omics studies. This beetle shows a more insect-typical development compared to flies, and has an extensive genetic toolkit that allows, for instance, the transgenic marking of neural cell types and in-depth gene function studies. Concerning the amenability to research and the extent of the toolkit, *Tribolium* is second only to *Drosophila,* offering a balance of evolutionary significance and experimental practicality (Berghammer et al., 1999; Campbell et al., 2022; Dönitz et al., 2018; Farnworth et al., 2020a; Gilles et al., 2015; Hakeemi et al., 2022; Herndon et al., 2020; Irion and Nusslein-Volhard, 2022; Klingler and Bucher, 2022; Mau et al., 2023; Strobl et al., 2015).

As proof of principle, we present two methods to dissociate and sort labelled neural cells and nucleus suspensions for single cell transcriptomics in two developmental stages of *Tribolium.* For the first strategy, we chose the neural stem cells, also named neuroblasts (NBs). These cells delaminate from the neuroectoderm during embryogenesis to generate two main cell types: Type I NBs and Type II NBs. They divide asymmetrically and produce intermediate progenitors that are in charge of generating the neural cell diversity of the insect central nervous system (CNS) (Arefin et al., 2020; Biffar and Stollewerk, 2014; Hartenstein and Wodarz, 2013; Hartenstein et al., 2021; Li et al., 2013; Thor, 2024a; Urbach and Technau, 2003; Urbach et al., 2016; Walsh and Doe, 2017). In the second strategy, we focused on neurons expressing the *Retinal homeobox* (*Rx*) gene, a conserved transcription factor involved in brain development in *Tribolium* larvae. Progenitor cells expressing *Rx* give rise to key neuron lineages in higher-order structures such as the mushroom bodies and central complex. These neuropiles play significant roles in sensing and integrating information, establishing motor coordination, and processing learning (Farnworth et al., 2020b). In both cases, neuroblasts and *Rx*-neurons, comparative work has revealed conserved and diverged aspects, but single-cell comparative studies are still missing. In summary, given that the knowledge about neural cells is restricted to the model organism *Drosophila*, a comparative single-cell sequencing approach using *Tribolium* is very promising.

Single-cell studies require high-quality cell or nuclei suspensions as input, which are usually laborious to produce because they require a high degree of standardization and optimization. The successful generation of a high-quality dataset for single-cell sequencing relies on the integrity, yield, and purity of the suspension used, which is influenced by multiple factors. For example, the organism, tissue, experimental design, and even operator experience can influence the quality of the cell suspensions produced, making this step the most challenging and variable in single-cell workflows (Denisenko et al., 2020; Machado et al., 2021; McLaughlin et al., 2022b; Regan and Preall, 2022; Wiegleb et al., 2022). To address this bottleneck, we tailored two robust dissociation protocols and FACS-based sorting strategies for neural cell types of *Tribolium castaneum.* In the first approach, we applied our strategy to isolate and enrich a cell suspension with embryonic NBs from a transgenic reporter line. The result was a high-resolution single-cell transcriptomic dataset of embryonic NBs. Secondly, we devised a method to dissociate and enrich for nuclei from neuronal lineages expressing the gene *Rx* in the larval brain. Yielding an enhanced single-nuclei dataset specific to these neuronal lineages. Based on these results, we posit that these protocols are adaptable to other tissues, developmental stages, and other insect species, providing a significant framework for robust single-cell comparative studies of rare neural cell types in insects.

## Materials and Methods

The two detailed protocols for cell and nuclei dissociation and FACS sorting described in this work are also available in Protocols.io: [Workflow 1](https://www.protocols.io/private/665CFC86CE7A11EF96BB0A58A9FEAC02) and [Workflow 2](https://www.protocols.io/private/BE2C2ABCA5ED11F19BD80A58A9FEAC02).

### Workflow 1: embryonic neuroblasts

#### Animal husbandry and embryo collection

*Tribolium castaneum* strain San Bernardino was reared for experimental procedures in boxes with wheat flour type 1050 maintained at 32°C at 40% humidity. The beetles were transferred into boxes with white wheat flour type 450 at 32°C for 12 hours to collect embryos. The laid eggs were collected and aged until they reached the developmental embryonic stage required for the experiments.

#### Embryo fixation, RNA probes, and FISH

The embryo fixation, RNA probes, and fluorescent *in-situ* hybridization of the *Tribolium* embryos were performed following protocols previously described (Buescher et al., 2020).

#### Cell dissociation of *Tribolium* embryos

*Tribolium* embryos were transferred into a washing basket and submerged in Milli-Q water to remove as much flour as possible. We proceeded to dechorionate the embryos with 5% bleach for three minutes and removed all traces of bleach with Milli-Q water. We transferred the embryos into a 2 mL cell douncer (Merck D8938) prefilled with 1 mL of ice-cold modified Schneider’s medium (Schneider’s medium mixed with 1% BSA, Thermo Fisher Scientific 21720024 and 15260037) using a paintbrush and proceeded with the mechanical dissociation by approximately 15-20 strokes of the pestle A. The homogenate was decanted into a 1.5 mL microcentrifuge tube, and an extra 500 µL of modified Schneider’s medium was used to wash and collect any cells remaining in the douncer and decanted into the same tube. The vial was centrifuged for three minutes at 50g at 4°C, and the resulting supernatant was decanted into a new 1.5 mL tube that was centrifuged for 5 minutes at 300g at 4°C. At this point, we discarded the supernatant by decanting while carefully avoiding losing the pellet. Using a 1 mL pipette tip, we added 1 mL of modified Schneider’s medium and gently pipetted up and down as much as necessary to dissolve the pellet. We previously coated a 1.5 mL centrifuge tube with modified Schneider’s medium and placed a 20 μm cell strainer (PluriSelect 43-10020-60) on top of the tube to filter the cell suspension. Extra 500 μL of modified Schneider’s medium was used to fully collect the cell suspension from the centrifuge tube and the cell strainer. The filtered cell suspension was centrifuged for 5 minutes at 300g at 4°C, and the supernatant was discarded by decanting. Finally, we dissolved the pellet in 50 μL of resuspension buffer (Schneider’s medium, 1% BSA, 0.8% Protector RNase-Inhibitor (Roche 3335402001), 1x Protease Inhibitor Cocktail (Promega G6521). We maintained the resulting cell suspension on ice and immediately transported it to the sorting facilities.

#### FACS of embryo cell suspensions

Cell suspensions of *Tribolium* embryos were immediately processed using a cell sorter model SH800S (Sony Biotechnology Inc.) with a 100-μm microfluidic sorting chip. As controls, before sorting our sample, we always analysed cell suspensions of wild-type *Tribolium* embryos to ensure that our gating strategy could differentiate DAPI-negative cells and to ensure that no false GFP-positive cells were sorted. We then stained the sample with DAPI to a concentration of 700 ng/mL for 15 minutes. We started by analysing all events using the forward scatter (FSC-A) and the side scatter (BSC-A) without gating. We created a second plot of all events using forward scatter (FSC-A) and fluorescence (DAPI). In this plot, we used a gate to select DAPI-negative cells. These events were plotted again using phycoerythrin (PE) versus GFP to exclude auto-fluorescent cells, but also to identify a small population of highly GFP-fluorescent cells that we gated into a final plot using forward scatter (FSC-A) and side scatter (FSC-W) to discard doublets. We sorted the GFP cells into a centrifuge tube coated and prefilled with 50 μL of ice-cold resuspension buffer, aiming to perform the procedure in no more than an hour.

#### 10x single-cell sequencing of enriched neuroblasts

The sorted cells were processed using the Chromium Next GEM Single Cell 3ʹ Reagent Kits v3.1 and a 10x controller, attempting to recover 10,000 cells (Chromium Single Cell 3’ Reagent Kits User Guide (v3.1 Chemistry Dual Index) - Official 10x Genomics Support). We, however, loaded 25,000 cells instead of the recommended 16,000 cells. It has been reported that this is necessary when performing 10x single-cell with *Drosophila* cells because insect cells are generally smaller than mammalian cells (McLaughlin et al., 2022b; Wiegleb et al., 2022). Libraries produced were sequenced using an Illumina NextSeq 2000 System to an average depth of 50,000 reads per cell.

#### Single-cell data analyses of neuroblasts

To further process our single-cell datasets, we used Cell Ranger pipelines to perform barcode processing, read alignment against the *Tribolium castaneum* transcriptome (Tcas5.2) (Dönitz et al., 2015), gene counting, and sample aggregation. The resulting dataset was further analysed using Scanpy (Wolf et al., 2018). First, we checked the number of cells, genes, UMI counts, percentage of mitochondrial reads, and percentage of ribosomal reads to define the filtering thresholds that maintain only data of the best quality. Secondly, using Scanpy and Python scripts, we normalized, log-transformed, and performed dimensionality reduction analyses and clustering of the data.

### Workflow 2: Rx-neurons

#### Animal husbandry and larval collection

*Tribolium castaneum* strain San Bernardino was maintained on instant flour in a humidity-controlled incubator at 28°C. Adults were allowed to lay eggs for 24 h. Afterwards, beetles were sieved to separate embryos, which were transferred into 300-µm mesh inserts placed in six-well plates. Larvae were allowed to hatch and develop for a further 4 days at 28°C. Early first-instar (L1) larvae were used for brain dissections.

#### Larval brain dissection, fixation, and immunolabeling

Larval brain dissections were performed essentially as described previously (Hunnekuhl et al., 2020). Early L1 larvae were dissected in ice-cold PBS using Dumont 5SPSF forceps. The anterior region was separated from the trunk, and residual non-neural tissue was removed. Isolated brains were transferred to fresh black dissection dishes maintained on ice. Individual dissection sessions were limited to 1 h.

Brains were fixed in 1 mL 1× dithiobis(succinimidyl propionate) (DSP) fixation solution for 2 hrs on ice. DSP fixation solution was prepared by diluting 50× DSP stock (50 mg/ml) into PBS immediately before use. PBS was added dropwise while vortexing to minimize DSP precipitation (Attar et al., 2018). Fixed brains were rinsed once and washed three times for 5 min in PBT (PBS containing 0.3% Triton X-100) on ice. Brains were then washed three times for 5 min in PBS on ice and transferred to PBS supplemented with 0.1 U/µL Protector RNase Inhibitor. At this stage, fixed brains were stored at 4°C for up to 7 days until sufficient material had been collected. In total, approximately 650 larval brains were pooled for nuclei isolation. Brains were not dehydrated or stored in methanol because this impaired staining with the anti-Rx antibody.

For immunolabeling, brains were washed six times for 5 min in PBT on ice and blocked for 2 hrs at 4°C in blocking solution consisting of 5% bovine serum albumin (BSA) in PBS. Brains were incubated overnight at 4°C with guinea pig anti-Tc-Rx primary antibody (1:1,000; Farnworth et al., 2020) diluted in antibody solution (2% BSA and 0.25% Protector RNase Inhibitor in PBS). Samples were rinsed three times in PBT and washed three times for 30 min in PBT at 4°C. Brains were subsequently incubated overnight at 4°C with goat anti-guinea pig Alexa Fluor 647 secondary antibody (1:1,000; Thermo Fisher Scientific, A-21450). Following secondary-antibody incubation, brains were rinsed three times in PBT and washed three times for 30 min in PBS at 4°C. Immunolabeled brains were pooled in PBS containing 0.1 U/µL Protector RNase Inhibitor.

#### Nuclei isolation from fixed larval brains

All nuclei-isolation steps were conducted using pre-chilled buffers, with samples maintained on ice unless otherwise stated. Immunolabeled brains were briefly centrifuged to collect the tissue, and excess PBS was carefully removed. Brains were resuspended in 1 mL ice-cold homogenization buffer and transferred to a sterile 2-mL Dounce homogenizer.

Homogenization buffer consisted of nuclear isolation medium (NIM; 250 mM sucrose, 25 mM KCl, 5 mM MgCl22, and 10 mM Tris-HCl, pH 8.0) supplemented with 100 µM dithiothreitol (DTT), 1× protease inhibitor cocktail, 0.5% Protector RNase Inhibitor, and 0.1% Triton X-100 (Wiegleb et al., 2022). Brains were mechanically dissociated with approximately 20 strokes using the loose pestle, followed by 40 strokes using the tight pestle. Care was taken to avoid bubble formation. Homogenates were incubated on ice for 10 min and mixed by gentle pipetting to ensure complete tissue disruption.

The homogenate was transferred to a 1.5-mL low-binding microcentrifuge tube and centrifuged for 10 min at 500 × g and 4°C in a swing bucket centrifuge. The supernatant was removed without disturbing the nuclei pellet. Nuclei were resuspended in 1 mL resuspension buffer consisting of PBS supplemented with 2% BSA and 0.15% Protector RNase Inhibitor. The suspension was passed through a 20-µm cell strainer into a low-binding microcentrifuge tube. Tubes, strainers, and pipette tips were pre-coated with resuspension buffer to minimize nuclei loss. Filtered nuclei were centrifuged for 5 min at 500 × g and 4°C, and the pellet was resuspended in 600 µL ice-cold resuspension buffer. Nuclei suspensions were kept on ice and immediately processed for FACS.

#### Fluorescence-activated nuclei sorting of larval brain suspensions

Nuclei were sorted using an SH800S Cell Sorter fitted with a 100-µm microfluidic sorting chip. The sorter was calibrated before nuclei preparation, and sorting was performed at 4°C. Three control samples were used for spectral unmixing and gate definition: an unstained nuclei suspension, a DAPI-only control, and an Alexa Fluor 647-only control generated from Rx-immunostained brains.

Nuclei suspensions were stained with 6 µL of 100 µg/mL DAPI for 20 min on ice. Events were first visualized by forward scatter area (FSC-A) and side scatter area (SSC-A), and events in the lower quarter of the scatter plot were selected. Singlet nuclei were identified by the linear relationship between FSC-A and forward scatter height (FSC-H). DAPI-positive nuclei were then selected based on DAPI fluorescence versus SSC-A, using the unstained control to exclude DAPI-negative events. Finally, Alexa Fluor 647-positive nuclei were gated based on Alexa Fluor 647 fluorescence versus SSC-A; the DAPI-only control was used to define the Alexa Fluor 647-negative population. DAPI- and Alexa Fluor 647-double-positive nuclei were collected into low-binding microcentrifuge tubes pre-coated and prefilled with 200 µL ice-cold resuspension buffer.

To maximize recovery, sorting was continued until approximately 100 µL remained from the initial 600-µL nuclei suspension. An additional 300 µL resuspension buffer was added to the remaining sample, and sorting was continued to 100 µL. This dilution-and-sorting step was repeated once.

#### Reverse crosslinking and 10x Genomics library preparation of *Rx*-nuclei

DSP crosslinks were reversed after sorting to render nuclear RNA accessible for library preparation. Sorted nuclei in 200 µL resuspension buffer were supplemented with 750 µL resuspension buffer containing 50 mM DTT. Samples were incubated for 30 min at 37°C and centrifuged for 10 min at 500 × g at room temperature. Room-temperature centrifugation was used to prevent DTT precipitation. The supernatant was carefully removed, and nuclei were resuspended in 1 mL resuspension buffer, followed by a second centrifugation for 10 min at 500 × g. After removal of 940 µL supernatant, the nuclei pellet was gently resuspended in the remaining 60 µL resuspension buffer. Nuclei were counted, adjusted to the desired concentration, and immediately processed using the 10x Genomics Chromium single-nucleus RNA-sequencing workflow.

#### 10x single-cell sequencing of enriched *Rx*-nuclei

The sorted nuclei were processed using the GEM-X Universal 3’ Gene Expression Reagent Kits v4 and a Chromium iX controller, attempting to recover 5,000 cells (10x Genomics, Chromium GEM-X Single Cell 3’ v4 Gene Expression User Guide). Libraries produced were 150 bp paired-end sequenced using a NovaSeq X Plus System to an average depth of 50,000 reads per cell.

#### Single-nuclei data analyses of *Rx*-nuclei

To further process our single-nuclei dataset, we used Cell Ranger pipelines to perform barcode processing, reads alignment against the *Tribolium castaneum* transcriptome (Tcas5.2) (Dönitz et al., 2015), and gene counting. The resulting dataset was further analysed using Scanpy (Wolf et al., 2018). First, we checked for the number of nuclei, genes, UMI counts, percentage of mitochondrial reads, and percentage of ribosomal reads to define the filtering thresholds that maintain only data of the best quality. Secondly, using Scanpy and Python scripts, we normalized, log-transformed, and performed dimensionality reduction analyses and clustering of the data.

#### Image acquisition and analysis

We obtained confocal stack images with a Zeiss LSM 980 microscope and ZEN blue software. Final Images were organized with Adobe Photoshop (26.2.0), Affinity Designer 2 (2.2.1), and Adobe Illustrator CS5 (15.0.2).

## Results

In this work, we present *Tribolium* as a significant insect model for developmental comparative single-cell studies. In order to do this, we developed a comprehensive workflow addressing several key challenges in cell-targeted single-cell transcriptomic studies. First, we addressed the generation of two types of suspensions, one for live cells and another for nuclei. Second, we targeted two types of rare neural cell types: neuroblasts and *Rx*-neurons. Third, we profiled two developmental stages: embryos and larvae. Fourth, we implemented cell enrichment via FACS using two strategies: sorting of live cells based on endogenous fluorescence from a transgenic reporter line and sorting of nuclei based on antibody staining. Fifth, we validated the use of fixed and unfixed cells as input for single-cell sequencing. Sixth, we performed our experiments in a coleopteran insect, which opens the opportunity for comparative studies, especially between two distant insect orders, Coleoptera and Diptera.

To make our approach clear and describe the challenges we overcame, the description of our workflow is divided into two parts. In workflow 1, we described the generation of cell suspensions from a transgenic line that fluorescently labelled the embryonic neuroblasts and the use of FACS for cell enrichment. In workflow 2, we present the generation of nuclei suspensions from larval brains, where *Rx*-neurons were labelled with an antibody and enriched with FACS. Both enriched suspensions were used for the generation of high-quality single cell transcriptomic data of *Tribolium* (**Fig. 1**).

**Fig. 1.**
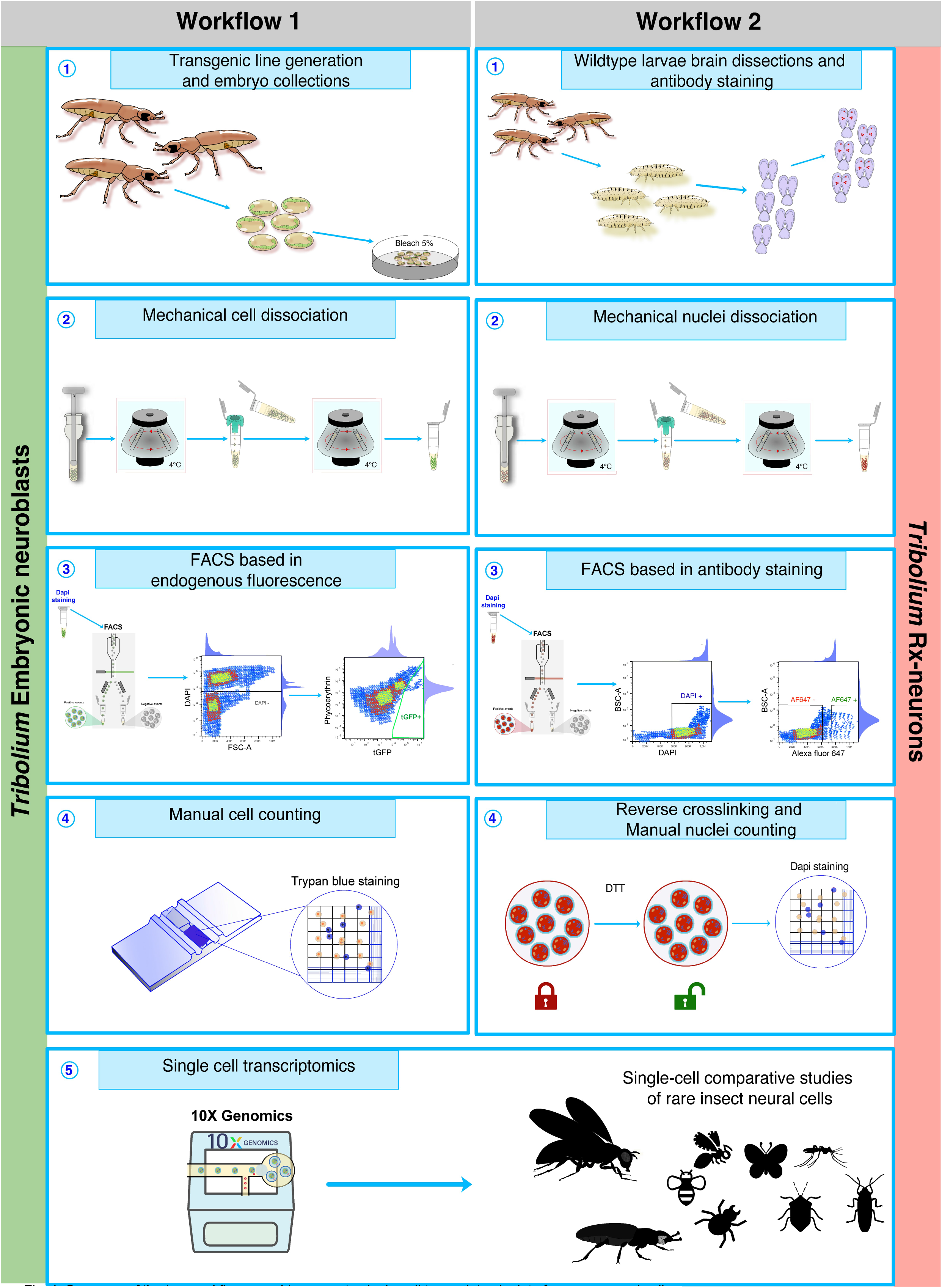
Summary of the two workflows used to generate single-cell transcriptomic data from rare neural cells. In workflow 1, we used a transgenic line to label embryonic neuroblasts **(1)**. We collected laid eggs and dechorionated them before proceeding to perform mechanical cell dissociation, which produced live cell suspensions **(2)**. We then stained the suspensions with DAPI before starting the cell sorting. DAPI was used to discard debris and dead cells, and endogenous green fluorescence was used to select the cells of interest **(3)**. Manual cell counting and inspection of fluorescence were performed under the microscope **(4)** before proceeding to perform single cell RNA sequencing **(5)**. In the second workflow, we collected wildtype larvae from which brains were dissected, fixed with DSP, and stained with an Rx antibody **(1)**. Mechanical nuclei suspensions were produced from the antibody-stained brains **(2)**. Nuclei suspensions were stained with DAPI, and the sample was sorted. DAPI was used to identify nuclei from debris, and the antibody fluorescence was used to enrich for Rx nuclei **(3)**. DSP fixation introduced crosslinks that were reversed using DTT **(4)**. Nuclei cell counting was performed before running single nuclei RNA experiments **(5)**.

### Workflow 1: embryonic neuroblasts

#### Defining the cells of interest during *Tribolium* embryogenesis

The insect CNS arises from embryonic neural stem cells that delaminate from the neuroectoderm and divide to generate the neural cell diversity of the VNC and brain. Despite being of key importance for CNS development, neuroblasts have been mainly studied in *Drosophila*, but their developmental dynamics are less known in other insects, including *Tribolium* (Biffar and Stollewerk, 2014; Biffar and Stollewerk, 2015; Urbach and Technau, 2003; Urbach et al., 2016). Therefore, to determine the developmental stages for sequencing, we identified the NBs across *Tribolium* embryonic development using the *Tc-asense* (*Tc-ase*) as a marker gene, which is the best-known marker for NBs in *Tribolium* (Biffar and Stollewerk, 2014; Wheeler et al., 2003). Fluorescent RNA *in-situ* hybridization experiments show that NBs are not present during early embryogenesis (NS0-2), and they only start appearing after NS3. Also, *Tc-ase* expression starts to decrease after NS14 (50 hours), which is in accordance with the NBs going into apoptosis or a quiescent state at the end of embryogenesis (Lai and Doe, 2014; Open et al., 2022). Since our goal was to obtain a comprehensive collection of NBs, we selected embryos between NS9-13 (30-50 hours at 32°C) as the ones that display the highest number of NBs and the highest *Tc-ase* expression (**Fig. 2A)**.

**Fig. 2.**
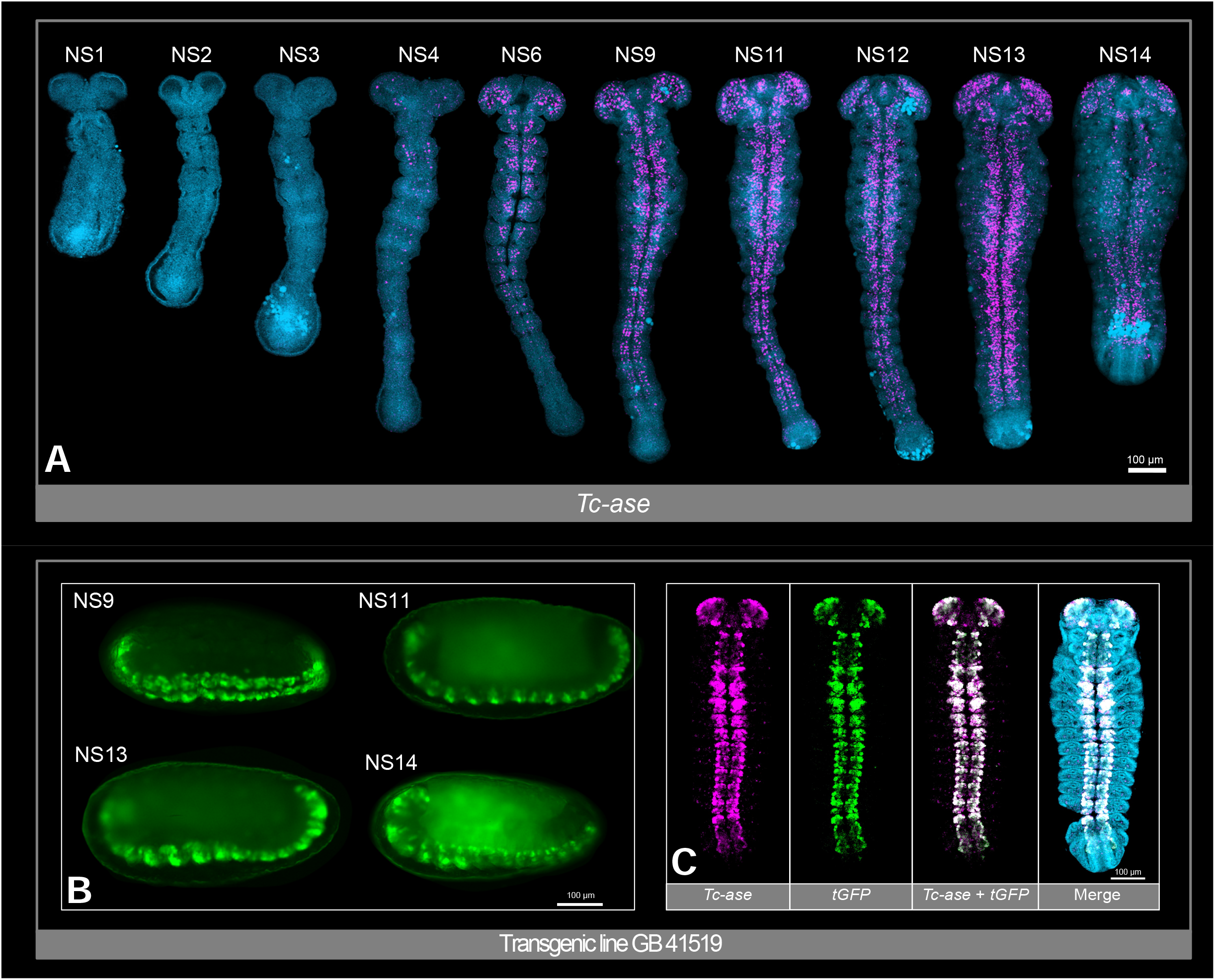
Tc-asense expression during *Tribolium castaneum* embryogenesis and characterization of the GB41519 reporter line labelling neuroblasts. **(A)** FISH of the *Tc-ase* gene in whole-mounted embryos reflects the NBs appearance dynamics. We show ten representative stages of *Tribolium* embryogenesis that correspond to neural stages previously described (Biffar and Stollewerk, 2014). We identified NS9 to NS13 (30-50 hours at 32°C) as the stages where the VNC and the brain display the highest number of NBs and the highest *Tc-ase* expression in *Tribolium* embryos. We chose these stages to enrich the cell suspensions with NBs. Nuclei are stained with DAPI (blue), and *Tc-ase* is stained in purple. **(B)** Endogenous fluorescence from the transgenic reporter line GB41519 that labels embryonic neuroblasts with a green fluorescent protein at several embryonic stages. **(C)** FISH detected the expression patterns of the gene *Tc-ase* (purple) and the tGFP transgene (green). We found that tGFP and *Tc-ase* expression patterns are very similar and largely overlapping (white). The merged figure shows the overlapping expression of *Tc-ase,* tGFP, and DAPI staining to reveal the shape of the embryo. Based on this analysis, we considered the cells marked by tGFP to be NBs. After cell dissociation, we performed cell sorting based on the tGFP signal of the NBs. Scale bars: 100 μm.

#### Using a transgenic reporter line to label cells

In *Drosophila,* we previously performed single-cell sequencing of the NBs using a *worniou* antibody (Cabañas et al., 2026), a protein found in the nuclei of NBs (Lai et al., 2012). However, this strategy is not feasible for *Tribolium* since there is no *worniou* antibody or any other antibody to mark the NBs. Therefore, we developed an alternative strategy by using a transgenic line in which a Gal4-enhancer trap drives UAS-tGFP (GB41519) in NBs (**Fig. 2B**) (Cabañas et al., 2026). To confirm the specific expression of the fluorescent protein in NBs, we performed a double staining of tGFP and *Tc-ase* (**Fig. 2C**). As this transgenic line expressed a cytoplasmic fluorescent protein, we performed single-cell rather than single-nuclei sequencing.

#### Cell dissociation and FACS sorting to enrich the labelled cells

We collected *Tribolium* embryos from the transgenic line and aged them until they were between 30-50 hours old. We removed the excess flour and dechorionated the embryos with bleach, then we thoroughly washed the embryos with water. We proceeded with the mechanical dissociation by placing approximately 200 μL of the embryos in a cell douncer prefilled with Schneider’s medium and using a pestle to disrupt the embryos. Afterwards, the sample was centrifuged, passed through a cell strainer, and centrifuged again (**Fig. 1. Workflow 1**). The resulting cell pellet was resuspended in Schneider’s medium with 1% BSA (**Additional file 1: Fig. S1. A and A’)**. The final sample was placed on ice and taken immediately to the sorting facilities, where we started by staining the cell suspension from our transgenic line with DAPI and used a second unstained cell suspension from wildtype *Tribolium* embryos to set the gating parameters. The unstained wildtype sample allowed us to have a control to reduce the amount of auto-fluorescent and false-positive cells (**Fig. 3**). After 15 minutes, our DAPI-stained sample was sorted to recover the GFP-positive cells (**Fig. 3. B, C, E,** and **F**). Our cell suspensions contained approximately 2×10^6^ cells in a 50 μL total volume (40×10^6^ cells/mL), which were sorted for no more than an hour to collect approximately 1-2,000 DAPI-negative and highly tGFP-positive cells (**Fig. 3E-F)**. To confirm the successful collection of tGFP-positive cells and quantify their yield, we inspected the cell suspensions before and after cell sorting to make sure that endogenous fluorescence was present (**Fig. 3. G, G’, H,** and **H’**). Finally, we performed manual cell counting using a Neubauer chamber (Kumar et al., 2019). We also tested whether this method could be scaled up, as single-cell methods generally require large numbers of input cells and biological replicates. The same cell dissociation protocol can be scaled up to obtain 20-40 ×10^3^ GFP-positive cells (**Additional file 1: Fig. S2. A)**.

**Fig. 3.**
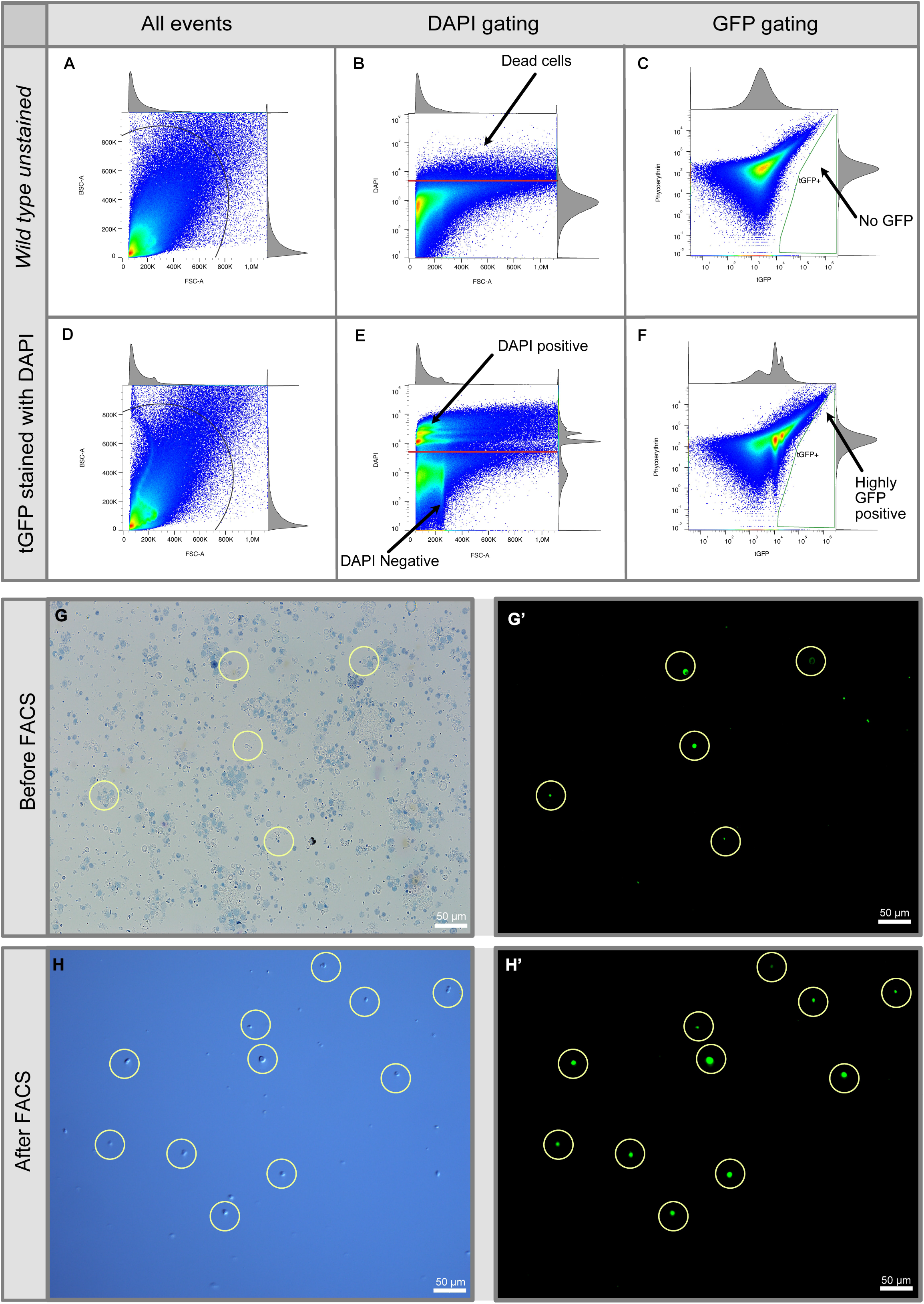
Methodology used to produce cell suspensions and expected results. Cell suspensions from wildtype were sorted to check all events **(A)**. As this suspension served as a control for setting gating parameters, cells were not stained with DAPI **(B)**, and they did not show high GFP fluorescence **(C)**. The cell suspension from the transgenic reporter line was processed in the same way. First, we identified all events **(D)**. This sample was stained with DAPI to exclude dead cells **(E)**. Finally, using the endogenous fluorescence, a small population of highly fluorescent cells was identified **(F)**. Phycoerythrin was used to exclude autofluorescent cells in both samples. The cell suspensions from the dissociation of embryos from a *Tribolium* transgenic line before and after sorting are shown in bright-field images and stained with trypan blue (**G** and **H**). The yellow circles show the endogenous fluorescence of the cells in the sample before and after sorting (**G’, H’**). Scale bars: 50 μm.

#### Single-cell RNA sequencing of *Tribolium* neuroblasts

The enriched cell suspensions were used for single-cell RNA sequencing with the 10X Genomics Chromium Technology (Chromium Single Cell 3’ Reagent Kits User Guide (v3.1 Chemistry Dual Index) - Official 10x Genomics Support). Libraries were sequenced using Illumina technology to an average depth of 50,000 reads per cell. We used ∼25-30,000 sorted cells for one run, resulting in ∼9,000 sequenced cells. FASTQ files were processed with the Cell Ranger pipeline to obtain matrices that contained the expression values. The datasets were further analysed using Scanpy (Wolf et al., 2018), and after quality control analyses, we recovered an average of 5,000 cells per sample. We confirmed that the tGFP protein was expressed in 80 percent of the sequenced cells, confirming successful FACS enrichment. To ensure that the information from our cells was reliable, we only kept cells that had more than 1,000 reads and less than 50,000 reads. We also restricted the number of mitochondrial reads to 25%, and the minimum number of genes that a cell should present was set to 500, and genes had to be present in at least 10 cells to be considered in our analyses (**Fig. 4)**. This workflow has already been used to produce a large, high-quality single-cell dataset from *Tribolium* neuroblasts in our recently released work (Cabañas et al., 2026). We also show, as an example, a subset of the data obtained. These cells are presented in UMAP plots that show unsupervised clustering (**Fig. 4A)**, expression of the green fluorescent protein used to label the neuroblasts (**Fig. 4B),** the total counts of each cell (**Fig. 4C)**, the number of genes captured in each sequenced cell (**Fig. 4D)**, and the percentage of mitochondrial reads present in each cell (**Fig. 4E)**. These plots confirm that we were able to obtain high-quality single cell data using our workflow.

**Fig. 4.**
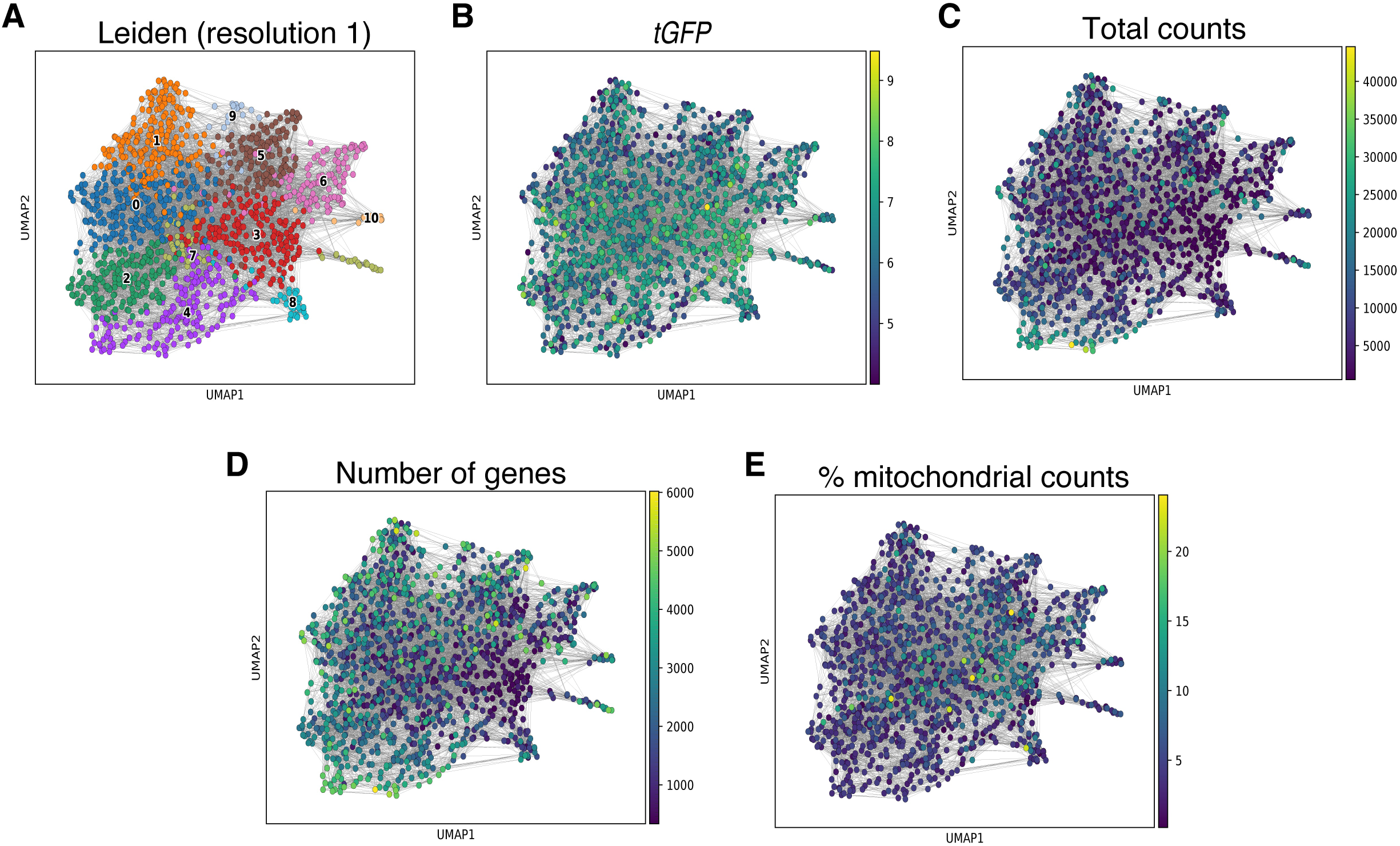
UMAP plots showing high-quality transcriptome obtained with workflow 1. A UMAP plot of unsupervised clustering of a subset of the cells sequenced **(A)**. Confirmation of the transgene *tGFP* used to label the neuroblasts sequenced **(B)**. We also evaluated the total counts captured **(C)**, the number of genes present in each cell **(D)**, and the percentage of mitochondrial reads in each cell **(E)**.

### Workflow 2: *Rx*-neurons

#### Antibody-based labelling enables enrichment of specific larval brain nuclei

The isolation of defined neural cell populations from the postembryonic *Tribolium castaneum* brain is challenging because available transgenic reporters are often cytoplasmic and therefore cannot readily be used to identify isolated nuclei. In addition, the larval brain is less amenable to rapid live-cell dissociation than embryos. We therefore established an antibody-based single-nucleus RNA-sequencing (snRNA-seq) workflow in which fixed larval brains were immunolabelled before nuclei isolation and FACS.

As a proof of principle, we used an antibody against retinal homeobox (*Tc-Rx*), a nuclear transcription factor expressed in mushroom body neuroblasts (MBNBs) and cells of the central complex (CX) in *Tribolium* (Farnworth et al., 2020b)(**Fig. 5**). Because *Tc-Rx* is localized to the nucleus, antibody labelling could be retained after tissue homogenization and used directly to identify target nuclei by FACS. This strategy therefore provides an alternative to transgenic fluorescent reporters and allows nuclei from fixed larval brains to be enriched for transcriptomic analysis. The approach is particularly useful for cell populations for which no suitable transgenic line is available or for experiments requiring the pooling of tissue collected across several dissection and fixation sessions.

**Fig. 5.**
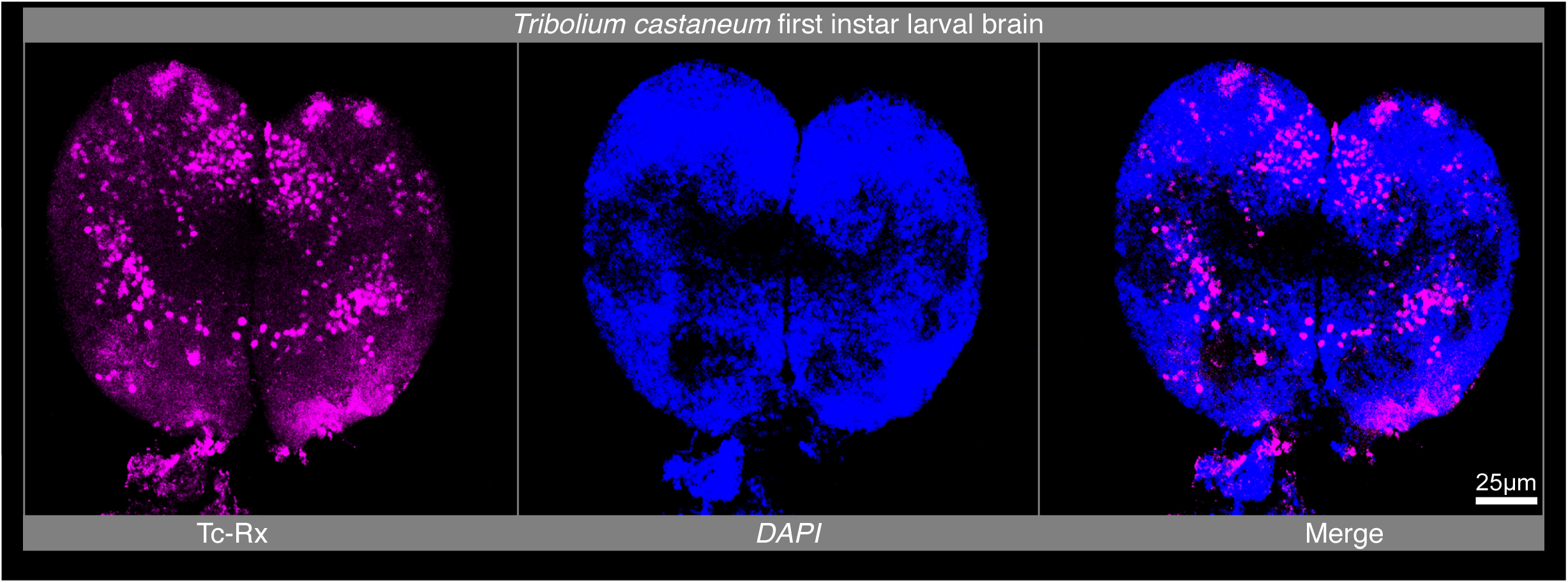
Tc-Rx immunolabelling in the early larval *Tribolium castaneum* brain. Maximum-intensity projection of an early first-instar larval brain stained with an antibody against *Tribolium* Retinal homeobox (Tc-Rx; magenta) and DAPI to label nuclei (blue). Tc-Rx-positive nuclei are detected in a restricted subset of cells, including the mushroom body neuroblasts and cells of the central complex. Scale bar: 25 µm.

#### Fixation and immunostaining preserve *Tc-Rx*-positive larval brains for nuclei isolation

We dissected early first-instar *Tribolium* larval brains, fixed them with the reversible crosslinker DSP, and pooled approximately 650 brains for immunolabelling and nuclei isolation. DSP fixation enabled tissue collection over several preparation rounds while retaining the Tc-Rx antibody signal. Because methanol storage interfered with Tc-Rx immunostaining, fixed brains were instead stored in PBS supplemented with RNase inhibitor until enough brains were fixed. Fixed brains were stained with a guinea pig anti-Tc-Rx antibody (Farnworth et al., 2020b) and an Alexa Fluor 647-conjugated secondary antibody. As expected from the restricted expression of *Tc-Rx* in MBNBs and CX cells, only approximately 5% of recovered nuclei were Alexa Fluor 647-positive (**Fig. 6F**). We mechanically dissociated the immunostained brains and obtained suspensions containing predominantly round, intact DAPI-positive nuclei (**Fig. 6G**). Alexa Fluor 647 fluorescence was retained in a subset of nuclei after dissociation (**Fig. 6G’+G’’**), confirming that Tc-Rx-positive nuclei could be identified and enriched by FACS.

**Fig. 6.**
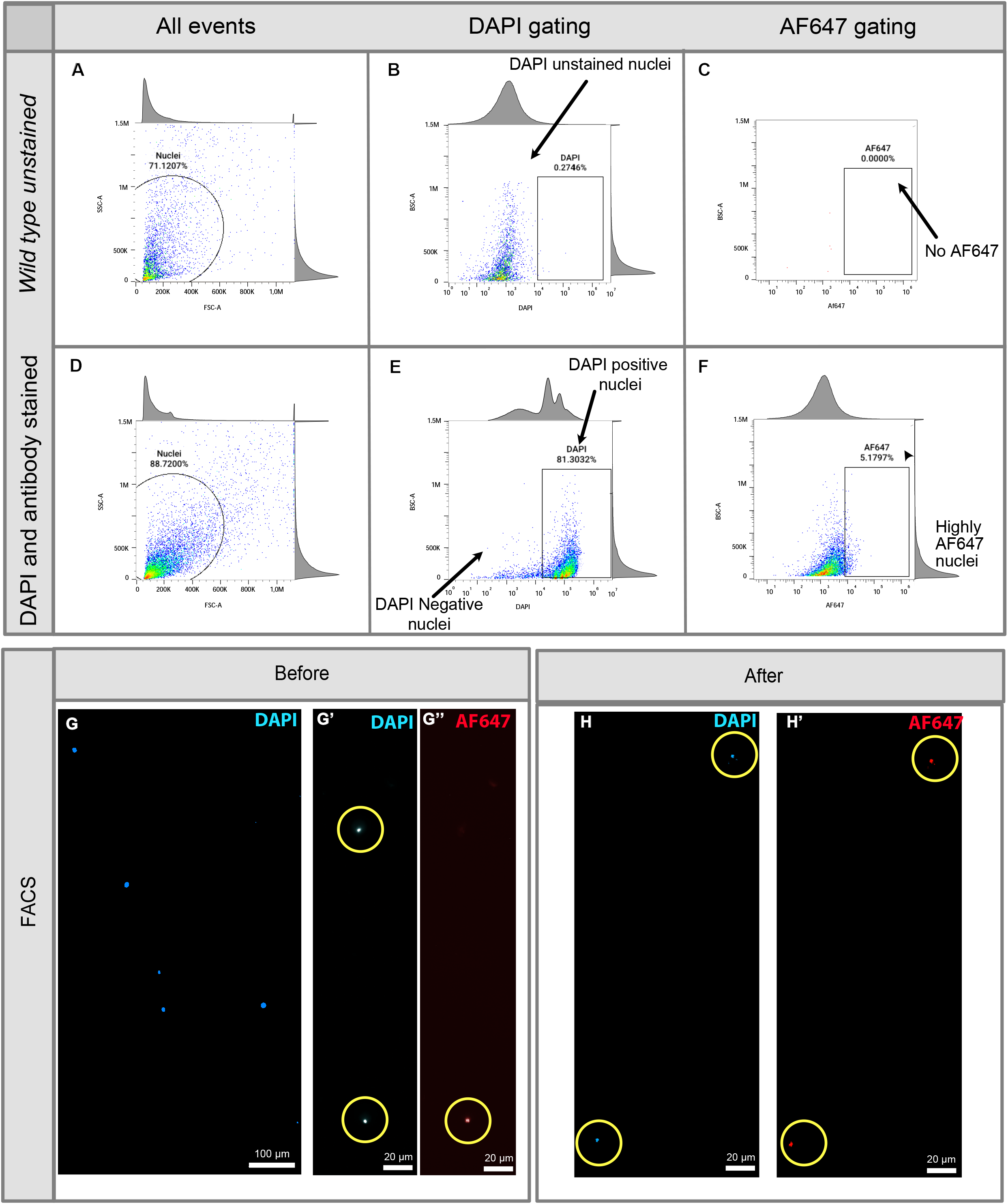
Fluorescence-activated sorting of Tc-Rx-labelled larval brain nuclei and expected results. Nuclei suspensions from unstained wild-type *Tribolium castaneum* larval brains served as controls for setting the gating parameters. First, nuclei were identified based on their scatter properties (**A**). These unstained nuclei showed no substantial DAPI (**B**) or Alexa Fluor 647 (AF647) fluorescence (**C**). Nuclei suspensions from DAPI-stained, Tc-Rx-immunolabelled larval brains were processed using the same gating strategy. First, nuclei were identified based on their scatter properties (**D**). DAPI-positive nuclei were selected to exclude DAPI-negative events (**E**), and a population with high AF647 fluorescence was identified for sorting (**F**). Representative fluorescence images show nuclei before (**G**–**G″**) and after sorting (**H**, **H′**). Before sorting, DAPI staining identified nuclei in the suspension (**G**, **G′**), whereas AF647 fluorescence was detected in a subset of nuclei (**G″**). After sorting, recovered nuclei showed both DAPI (**H**) and AF647 fluorescence (**H′**). Yellow circles indicate corresponding nuclei in the DAPI and AF647 channels. Scale bars: 100 μm in G; 20 μm in G′, G″, H and H′.

#### FACS enriches Tc-Rx-positive nuclei

Nuclei suspensions were stained with DAPI before sorting. An unstained control was used to optimize the gating strategy and to define the DAPI- and Alexa Fluor 647-positive populations (**Fig. 6A-C**). After exclusion of debris and doublets, DAPI-positive nuclei were gated for Alexa Fluor 647 fluorescence to selectively recover Tc-Rx-positive nuclei (**Fig. 6D-F**). From approximately 650 larval brains, we recovered approximately 8,000 DAPI- and Alexa Fluor 647-double-positive nuclei, which were collected for downstream processing. Because DSP fixation introduces reversible crosslinks, we performed DTT-mediated reverse crosslinking after FACS sorting. Nuclei were inspected under the microscope (**Fig. 6H+H’**) and resuspended at the desired concentration for immediate further processing.

#### Single-nuclei RNA sequencing of *Rx*-positive *Tribolium* larval brain nuclei

The enriched nuclei suspensions were used for single-nuclei RNA sequencing with the latest 10X Genomics Chromium Technology for 3’ gene expression analysis (10x Genomics, Chromium GEM-X Single Cell 3’ v4 Gene Expression User Guide). Libraries were 150 bp paired-end sequenced using Illumina technology to an average depth of 50,000 reads per nucleus. We used ∼8,000 sorted cells for one run, resulting in ∼2,000 sequenced nuclei. FASTQ files were processed with the Cell Ranger pipeline to obtain matrices that contained the expression values. The datasets were further analysed using Scanpy (Wolf et al., 2018), and after the quality control analysis, we recovered an average of 1,000 cells per sample. We detected *Rx* transcripts in approximately 30% of the sequenced nuclei, confirming enrichment of the Rx-positive population by antibody-based FACS. The higher proportion of nuclei recovered by Tc-Rx immunolabelling than expressing detectable *Rx* transcripts is consistent with the persistence of Tc-Rx protein after *Rx* transcription has ceased. Thus, the antibody-based strategy also captured nuclei derived from cells with prior *Rx* expression, in addition to cells actively transcribing *Rx* at the time of collection. To ensure that the information from our nuclei was reliable, we only kept nuclei that had more than 1,000 reads and less than 50,000 reads. We also restricted the number of mitochondrial reads to 25%, and the minimum number of genes that a nucleus should present was set to 500, and genes had to be present in at least 10 nuclei to be considered in our analyses (**Fig. 7**). We show a subset of the cells we sequenced using our workflow 2 and processed according to the parameters we previously mentioned. In **Fig. 7A**, we show unsupervised clustering of the cells in our dataset. We also confirmed the expression of the *rx* transcript in these cells (**Fig. 7B**) and show the total number of counts per cell (**Fig. 7C**), the number of genes in each cell (**Fig. 7D**), and the percentage of mitochondrial reads in each cell (**Fig. 7E**). Apart from the *Tribolium* larval brain, we have also used this method to produce embryonic Neuroblasts using a worniou antibody in *Drosophila* (Cabañas et al., 2026).

**Fig. 7.**
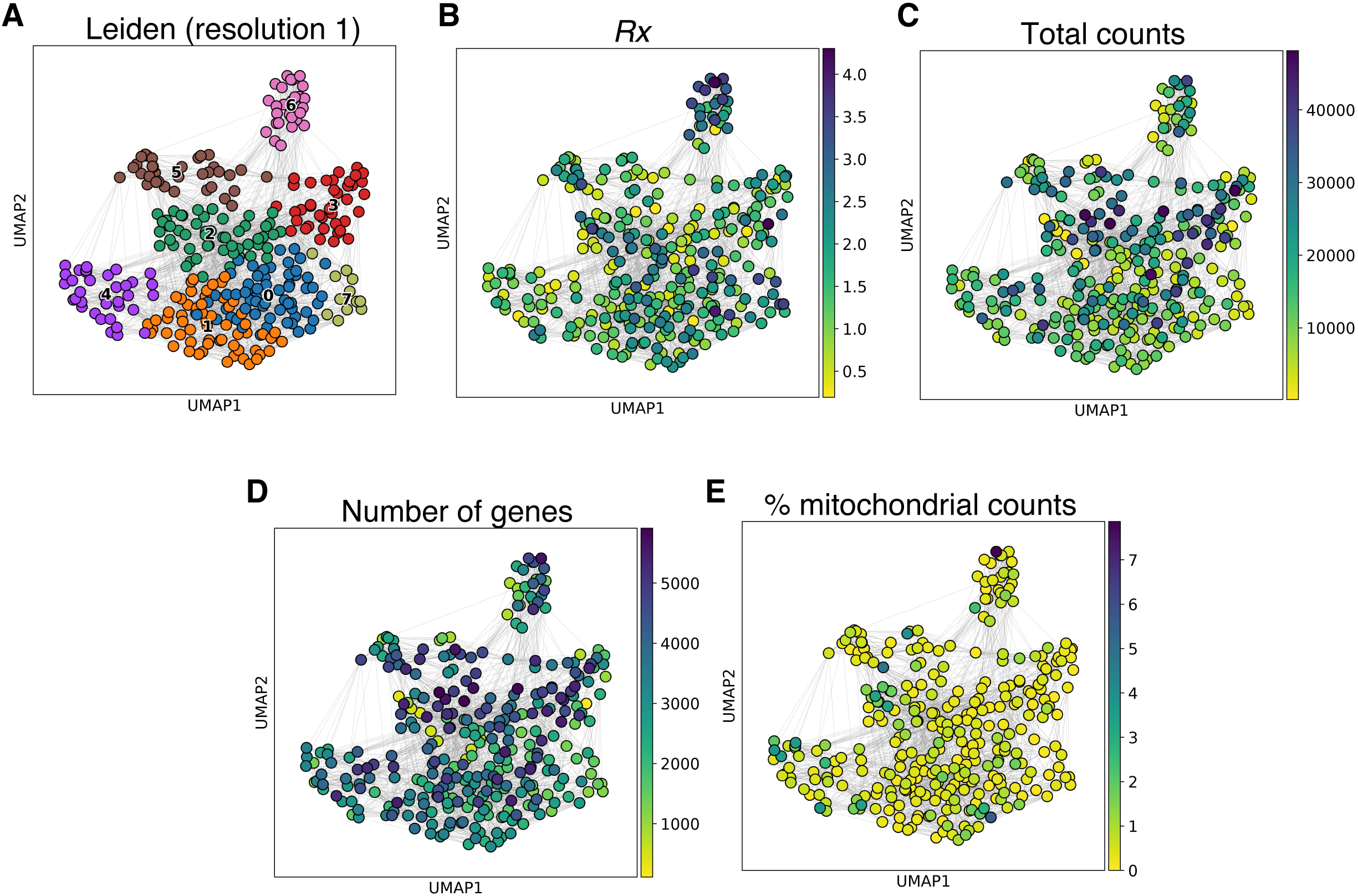
UMAP plots showing high-quality transcriptome obtained with workflow 2. UMAP visualization of unsupervised clustering of a subset of the sequenced nuclei (**A**). Detection of rx transcripts in the nuclei isolated using the Rx antibody (**B**). Total transcript counts per nucleus (**C**), number of detected genes per nucleus (**D**), and percentage of mitochondrial reads per nucleus (**E**).

## Discussion

In this work, we show how to obtain high-quality cell suspensions enriched with tGFP-labelled cells from embryos of *Tribolium* to produce single-cell RNA sequencing data. We further demonstrate that this workflow can be complemented by an antibody-based strategy for single-nuclei RNA sequencing, in which fixed larval brains are immunolabelled for the nuclear transcription factor *Tc-Rx* before nuclei isolation and FACS enrichment. This approach expands cell-targeted transcriptomic profiling in *Tribolium* beyond the use of available transgenic reporters and enables isolation of defined nuclear populations from postembryonic tissues. Below, we focus on factors that, based on our experience, have a significant effect when performing such experiments.

### Reducing cell stress is key when producing live cell suspensions

It has been reported that the quality of the cell suspension has a big impact on the resulting single-cell data (García-Castro et al., 2021; Marsh et al., 2022; Neuschulz et al., 2023; Piwecka et al., 2023; Wiegleb et al., 2022). Cells taken out of their homeostatic environment undergo considerable stress, increased by the enzymatic or mechanical steps that are necessary to obtain the dissociated cells. Such stress can generate aberrant expression patterns (Marsh et al., 2022; Van Den Brink et al., 2017). Therefore, reducing stress throughout the cell dissociation process is key to maintaining normal patterns of expression and keeping the cells viable during the whole procedure. We suggest avoiding the use of hazardous chemicals when treating animals or cells. For example, we used highly diluted 5% bleach in the dechorionate *Tribolium* embryos to avoid bleach-induced stress. We also chose Schneider’s medium for producing the cell suspensions since this has been widely used when working with insect cell cultures. Initially, we used PBS to perform the dissociation and obtained cleaner cell suspensions compared to Schneider’s medium. However, we realized that Schneider’s medium increased the number of live cells collected during cell sorting. We also realized that fetal bovine serum (FBS) increased cell viability, but also cell aggregation and amyloid-like aggregates. These observations are in accordance with studies that report the formation of cell aggregates of dissociated cells and cell cultures when media are supplemented with FBS (Heger et al., 2018; Lehrich et al., 2021; Matak et al., 2017). Therefore, we used bovine serum albumin (BSA) to obtain cleaner cell suspensions. It has also been reported that FBS can induce cell reprogramming, immune responses, and changes in gene expression levels (Kwon et al., 2016; Lehrich et al., 2021; Matak et al., 2017; Yamamoto et al., 2018). Moreover, we noticed that BSA and FBA at high concentrations (>10%) and mixed with Schneider’s medium increased the number of live cells recovered, but it has also been reported that high concentrations can interfere with the proper processing of the samples with the Chromium Next GEM Single Cell 3ʹ Reagent Kits (Chromium Single Cell 3’ Reagent Kits User Guide (v3.1 Chemistry Dual Index) - Official 10x Genomics Support). To balance these issues, we used 1% BSA in Schneider’s medium. This mix was also used for coating, as a cushion for cells, tubes, pipette tips, and cell strainers in order to reduce cell stress and increase the number of living cells recovered.

Sample temperature and processing time greatly impacted the outcome and should be carefully considered when preparing cell suspensions (Denisenko et al., 2020; Wiegleb et al., 2022). For testing purposes, we performed cell dissociation and sorting at room temperature conditions (**Additional file 1: Fig. S3**), and we were unable to collect any viable cells. So, it was necessary to always keep samples and solutions on ice and use the centrifuge at 4°C during the cell dissociation. Samples were always transported on ice, and sorting had to be performed at 4°C as well. Even with careful treatment, the cells are under substantial stress and might die quickly. Therefore, sample handling times were kept to a minimum. For instance, we spent 2 hours and 30 minutes from egg collection to cell suspension and performed FACS sorting within another 90 minutes. After quality control, we immediately ran our cell suspension in the 10x Genomics controller for GEM generation and barcoding library, and we subsequently performed GEM-RT cleanup and cDNA synthesis (Chromium Single Cell 3’ Reagent Kits User Guide (v3.1 Chemistry Dual Index) - Official 10x Genomics Support). Such a tight schedule depends on having well-trained personnel for each step of the methodology, or a team properly coordinated and trained to handle the beetles, perform the cell dissociation, carry out cell sorting, and conduct sequencing.

### Contamination and sample degradation can jeopardize sample quality

RNA degradation is a major concern during the preparation of cell and nuclei suspensions for single-cell and single-nucleus RNA sequencing. This is particularly relevant in workflows involving tissue dissociation, multiple wash steps, and prolonged immunolabeling incubations, during which RNA can be exposed to endogenous and environmental RNases. We therefore included Protector RNase inhibitor and a protease inhibitor cocktail throughout the sample-preparation workflows to protect RNA integrity and preserve cell/nuclei quality for the 10x technology (McLaughlin et al., 2022b; Sun et al., 2023; Wiegleb et al., 2022). The primary-antibody incubation requires particular attention because antisera and other animal-derived antibody preparations can contain RNase activity. Thus, we supplemented the antibody incubation buffer with RNase inhibitor to reduce the risk of RNA degradation during the overnight incubation of fixed larval brains with the Tc-Rx primary antibody. In our hands, the combination of one RNase inhibitor and one protease inhibitor cocktail was sufficient to obtain nuclei suspensions compatible with single-cell and single-nucleus RNA sequencing. While some protocols use several RNase inhibitors or higher inhibitor concentrations, adding a single RNase inhibitor at the relevant steps provided a practical balance between RNA protection, compatibility with immunostaining and downstream processing, and reagent costs. This is relevant when establishing targeted sequencing workflows in new tissues or organisms, where optimization often requires multiple preparative rounds and inhibitor costs can accumulate rapidly.

### Organism and cell type-specific requirements

Standardizing a method to produce high-quality enriched cell suspensions also depends on the organisms of choice. Hence, modifications will, in most cases, be necessary. For example, single-cell dissociation of dissected tissues from large models, like mice, is different compared to smaller organisms or small tissue samples (Huang et al., 2023; Wiegleb et al., 2022). The dissociation method will also be cell type dependent (Denisenko et al., 2020; Li, 2021; McLaughlin et al., 2022b; Wiegleb et al., 2022). For example, brain NBs are some of the biggest *Tribolium* cell types and are not part of an epithelium, which may predispose them to increased mechanical stress but ease dissociation. Our tests revealed that if centrifuged at high G-forces, the NBs die almost immediately, but if the centrifugation is not strong enough, the NBs are lost during the washing steps. We also tested filtering our cells using a 10 μm cell strainer, resulting in clean and uniform cell suspensions, but the number of GFP-positive cells recovered during sorting was reduced. On the other hand, using a 30 μm strainer allowed small chunks of tissue to go into the final cell suspension, causing clogging of the cell sorter. For NBs, we obtained the best results when using a 20 μm strainer (**Additional file 1: Fig. S1)**. Based on our experience, each step must be optimized extensively to make sure that live cells are recovered with high efficiency and minimal cell stress. The protocol presented here will have to be adapted with respect to these parameters in other cell types of the red flour beetle *Tribolium* and other insects with similar cell sizes.

### Single-nuclei sequencing versus single-cell sequencing

The single-nuclei sequencing method is able to endure longer sample handling times and is compatible with fixatives. Therefore, it is an excellent approach when samples need to be stored or are difficult to prepare, such that sample pooling is possible, and sorting over extended sorting periods could be an advantage too. In our case, when we attempted to use the *Drosophila* methods for producing nuclei suspensions, the fixation method did not work for *Tribolium* embryos. Second, we did not detect the cytoplasmic fluorescent protein in the nucleus after cell dissociation. This issue has been previously reported when dissociating *Drosophila* tissues (McLaughlin et al., 2022b). Therefore, we opted to produce single-cell suspensions of *Tribolium* embryonic neuroblasts, such that the cytoplasmic fluorescence of the cells was used for sorting. The downside of this single-cell sequencing approach was that cell viability, yield, quality, and cell clumping had to be considered, and the necessity to perform all steps of each preparation within a few hours.

### Using transgenic lines

Single-cell sequencing does not require transgenic animals or staining methods if all the cells of a tissue, an organ, or a complete animal are of interest. In these cases, however, the most abundant or the most resilient cells to the dissociation method will be enriched in the sample. Rare cell types will be underrepresented or neglected in the results. Also, sequencing depth is shared among all cells in the sample, so sequencing cost must be increased to maintain reliable sequencing coverage across the expanded number of profiled cells. On the other hand, sequencing a subset of isolated cells ensures that sequencing coverage focuses on a specific cell type, but this requires antibodies or a transgenic approach to mark the target cell population and the sorting of the samples.

As NBs are comparably rare in the embryo, enrichment was highly recommended. An antibody was used to mark and sort nuclei derived from NBs in flies (Cabañas et al., 2026). Since there is no antibody available for this purpose in *Tribolium*, we opted to use a transgenic line. The availability of transgenic lines may be a bottleneck and has been used comparably rarely (Huang et al., 2023; May and Röper, 2024; Secchia et al., 2022; Veloso, 2022; Velten et al., 2022; Vogler et al., 2021). Actually, it has not been previously used in emerging model systems because of the lack of their respective transgenic lines. The red flour beetle *Tribolium castaneum* sticks out in that respect, as there are a comparably large number of enhancer trap lines available (He et al., 2019; Trauner et al., 2009), and genome editing has recently been used to generate enhancer traps and bicistronic imaging lines (Bullinger, 2024; Farnworth et al., 2020a; He et al., 2019; Rethemeier et al., 2024a). Given the genome editing advancements in emerging model organisms, that approach might increase in importance in the future. Still, generating transgenic lines in emerging model organisms can be time-consuming, so labelling cells with cross-reacting antibodies might be an excellent alternative (Cabañas et al., 2026). As argued above, single nuclei sequencing has advantages, but the cytoplasmic location of the usual fluorescent proteins in existing imaging lines does not mark the nuclei well in our experience. Hence, nuclear localization of the fluorescent proteins should be considered. Lamin marked with a fluorescent protein has been reported to be suitable for sorting and single-cell sequencing in *Drosophila* (McLaughlin et al., 2022b), and we have tested in this work the use of the Tc-Rx antibody in *Tribolium*.

### Antibody-based enrichment for single-nuclei RNA sequencing

Antibody-based labelling provides a complementary strategy for enriching defined cell populations for single-nuclei RNA sequencing. Rather than relying on a transgenic reporter, this approach uses an antibody directed against a nuclear antigen that is selectively expressed in the cells of interest. Following nuclei isolation, antibody-positive nuclei can be identified by FACS and recovered for single-nuclei RNA sequencing. Antibody-based FACS of immunolabeled cells has been used to derive NBs in flies (Cabañas et al., 2026).

In our case, we used an antibody against Tc-Rx, a nuclear transcription factor expressed in MBNBs and CX cells. The nuclear localization of Tc-Rx was advantageous because the antibody signal remained associated with nuclei after dissociation. This allowed us to fix and immunolabel larval brains before nuclei isolation. Here we demonstrated that antibody-based enrichment can recover a defined and relatively rare larval brain population for downstream snRNA-seq.

An important advantage of this strategy is that it does not require the prior generation or availability of a transgenic line. This is particularly valuable in emerging model organisms, in which suitable cell-type-specific reporter lines may be unavailable, difficult to establish, or may label the target population with cytoplasmic fluorescent proteins that cannot be retained in isolated nuclei. By contrast, antibodies recognizing nuclear proteins can directly mark nuclei for sorting. The strategy can therefore extend snRNA-seq to enriched cell populations defined by the availability of a specific and robust antibody, including developmental stages or tissues for which live-cell dissociation is difficult.

Antibody-based enrichment also has practical advantages for the larval brain. In contrast to live-cell workflows, fixed nuclei can be collected over multiple dissection rounds and pooled before processing, reducing the pressure to complete dissection, dissociation, sorting, and library preparation within a single short time window. Reversible DSP fixation enabled us to collect larval brains over several days while preserving Tc-Rx immunoreactivity and subsequently reverse the crosslinks before 10x Genomics library preparation. Such fixation-compatible workflows can be especially useful for small tissues, low-abundance cell populations, or experiments requiring substantial starting material.

However, antibody-based sorting has limitations that should be considered during experimental design. First, enrichment depends on the specificity, sensitivity, and compatibility of the antibody with the chosen fixation and nuclei-isolation conditions. We found that methanol storage was incompatible with Tc-Rx immunostaining, illustrating that fixation and storage conditions need to be optimized individually for each antibody. Second, detection of a protein does not necessarily correspond to ongoing transcription of the respective gene. Indeed, *Rx* transcripts were detected in approximately 30% of sequenced nuclei, whereas all sorted nuclei were selected based on Tc-Rx antibody staining. This difference is consistent with persistence of Tc-Rx protein after *Rx* transcription has decreased or ceased. Therefore, antibody-positive nuclei should be interpreted as originating from cells with current or recent expression of the target protein, rather than exclusively from cells actively transcribing the corresponding gene at the time of dissection.

Finally, antibody incubations introduce additional handling time and potential sources of RNA degradation. RNase-free working conditions and the inclusion of RNase inhibitor in the relevant buffers, particularly during antibody incubation, are therefore important for maintaining RNA quality. Nonetheless, our results show that antibody-based nuclear labelling and FACS provide a feasible method for cell-specific single-nuclei RNA sequencing in *Tribolium*. Together with transgenic reporter-based live-cell sorting, this approach broadens the range of cell populations that can be selectively profiled during embryonic and postembryonic brain development.

### Novel cell enrichment methods not yet implemented in insects couple with single-cell sequencing

The tailored workflows presented in this work are versatile ways to advance the profiling of rare neural cell types in insects using enriched cell and nuclei suspensions coupled with single cell omics technologies. We expect the improvement of these methods and the addition of novel workflows. For example, recently several studies have taken advantage of hybridization chain reaction (HCR) RNA FISH to label rare cell populations and enrich the cell suspensions that are used to perform single cell transcriptomics (Abay et al., 2025; Amamoto et al., 2019; Chen et al., 2025; Liao et al., 2026; McKinney et al., 2026; Reilly et al., 2021). Our methods and recently devised HCR methods, coupled with sequencing that follow similar logic, are becoming a necessity for single cell studies in order to gain a deeper understanding of neural cells and how they compare across organisms. However, the HCR-based strategy has not been implemented in insect studies.

### Single-cell sequencing to study insect brain evolution and development

Single-cell sequencing is an excellent tool for performing single-cell comparative and cross-species single-cell studies for evolutionary-developmental questions. For example, *Drosophila* is the most widely used insect model organism for brain development in arthropods, and a huge body of knowledge has been accumulated on this topic (Thor, 2024b). However, it remains enigmatic to what extent the identified mechanisms diverge in other species. For example, it is known that there is variation in the number of brain NBs and the resulting brain morphology (el Jundi and Heinze, 2020; Rethemeier et al., 2024b; Urbach and Technau, 2003). For such questions, we propose *Tribolium castaneum* as a model for studying the diversification of neural development and its genetic underpinnings (Farnworth et al., 2020a; Farnworth et al., 2020b; Koniszewski et al., 2016; Rethemeier et al., 2024a).

## Conclusions

We established two complementary workflows for targeted transcriptomic profiling of rare neural populations in *Tribolium castaneum*: reporter-based enrichment of embryonic neuroblasts for single-cell RNA sequencing and antibody-based enrichment of Tc-Rx-positive larval brain nuclei for single-nucleus RNA sequencing. Together, these approaches extend access to selected neural populations across developmental stages, using either a transgenic fluorescent reporter or a nuclear antigen to identify the population of interest. Our results highlight the importance of adapting sample preparation to tissue properties, marker localization, and the requirements of downstream sequencing.

By enriching selected populations before sequencing, these workflows provide a means of investigating cells that may be underrepresented in whole-tissue or whole-organism transcriptomic surveys. They establish a methodological foundation for studying neural development in *Tribolium* and offer practical starting points for adaptation to other insect systems. This expands opportunities for comparative studies of neural cell diversity and the developmental mechanisms underlying insect brain evolution.

## Declarations

### Ethics approval and consent to participate

Not applicable.

### Consent for publication

Not applicable.

### Availability of data and materials

All relevant data and details of the resources used in this study are provided in the article and its supplementary information.

### Competing interests

The authors declare that they have no competing interests.

### Funding

This work was supported by the Deutsche Forschungsgemeinschaft (DFG; grant MA 10550/2-1 to CM).

### Authors’ contributions

CM, GB, and NC conceived the study. NC, CM, AV, and RZ performed the experiments. CM, GB, and NC prepared the manuscript. All authors read and approved the final manuscript.

## Acknowledgements

NC is a member of the International Max Planck Research School for Genome Science (IMPRS-GS) and gratefully acknowledges the support received. We thank Prof. Dr. André Fischer, Dr. Lalit Kaurani, and Susanne Burkhardt for providing facilities and supporting the sequencing experiments. We also thank Prof. Dr. Rüdiger Behr at the German Primate Center (DPZ) for providing access to the fluorescence-activated cell sorter.

## Additional files

File name: Additional file 1

File format: .pdf

Title of data: Supplementary figures on the optimization of embryonic cell suspension preparation in *Tribolium castaneum*.

Description of data: Supplementary figures S1–S3 showing the comparison of cell filtration conditions (Fig. S1), scaling up embryo dissociation for GFP-positive cell recovery (Fig. S2), and the effects of holding cell suspensions on ice or at room temperature for two hours (Fig. S3).

## Notes

https://www.protocols.io/private/665CFC86CE7A11EF96BB0A58A9FEAC02

https://www.protocols.io/private/BE2C2ABCA5ED11F19BD80A58A9FEAC02

### Competing Interest Statement

The authors have declared no competing interest.

