## Additional file 1 for "Single-cell transcriptomics of enriched rare neural lineages of *Tribolium castaneum*"

### Supplementary information

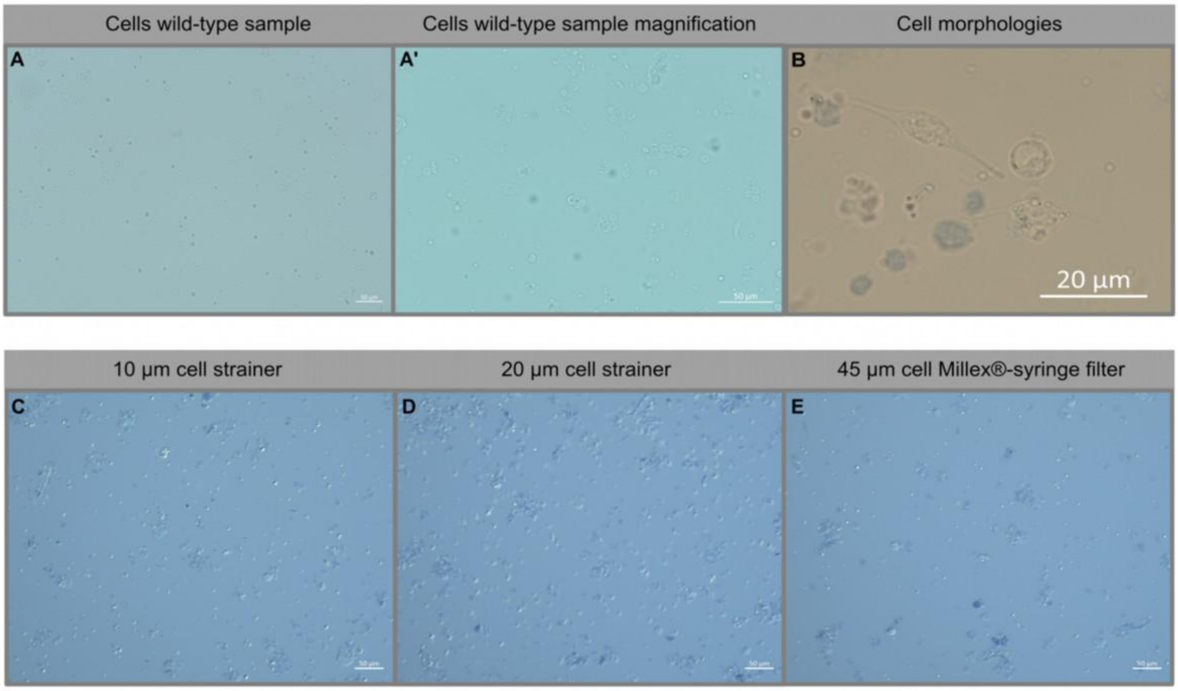


**Fig S1. Cell suspensions from wildtype *Tribolium* embryos.** In **Fig S1** (**A**), we show a cell suspension from *Tribolium* embryos with numerous cells. We show a magnification of the same sample in **Fig S1, A'**, where it is possible to identify cells of different sizes and how they can clump. Scale bar 50 µm. In **Fig. S1B**, we show that the cell suspensions from *Tribolium* embryos are not homogeneous and, as expected, multiple cell types are present. Here, we show examples of some of the morphologies we identified in our cell suspensions. When producing our cell suspensions, we tested and selected the best parameters. We show, as an example, the use of different cell strainers to remove chunks of tissue and to produce more uniform cell suspensions. The 10 µm cell strainer produced the most uniform and clean suspensions (**Fig S1, C**). However, the 20 µm strainer (**Fig S1, D**) allowed the recovery of more cells during the cell sorting compared to the 10 µm strainer and the 45 µm syringe filter (**Fig S1, E**). We also noticed that the 45 µm filter allowed the passing of big chunks of tissue into the final cell suspension, but it is still possible to recover GFP cells during the sorting from this suspension. We even recovered more cells from the suspension filtered with the 45 µm filter than with the 10 µm strainer. We obtained the best results, however, with the 20 µm strainer. All samples were stained with trypan blue to visualize cell viability in all samples. Scale bar 50 µm.

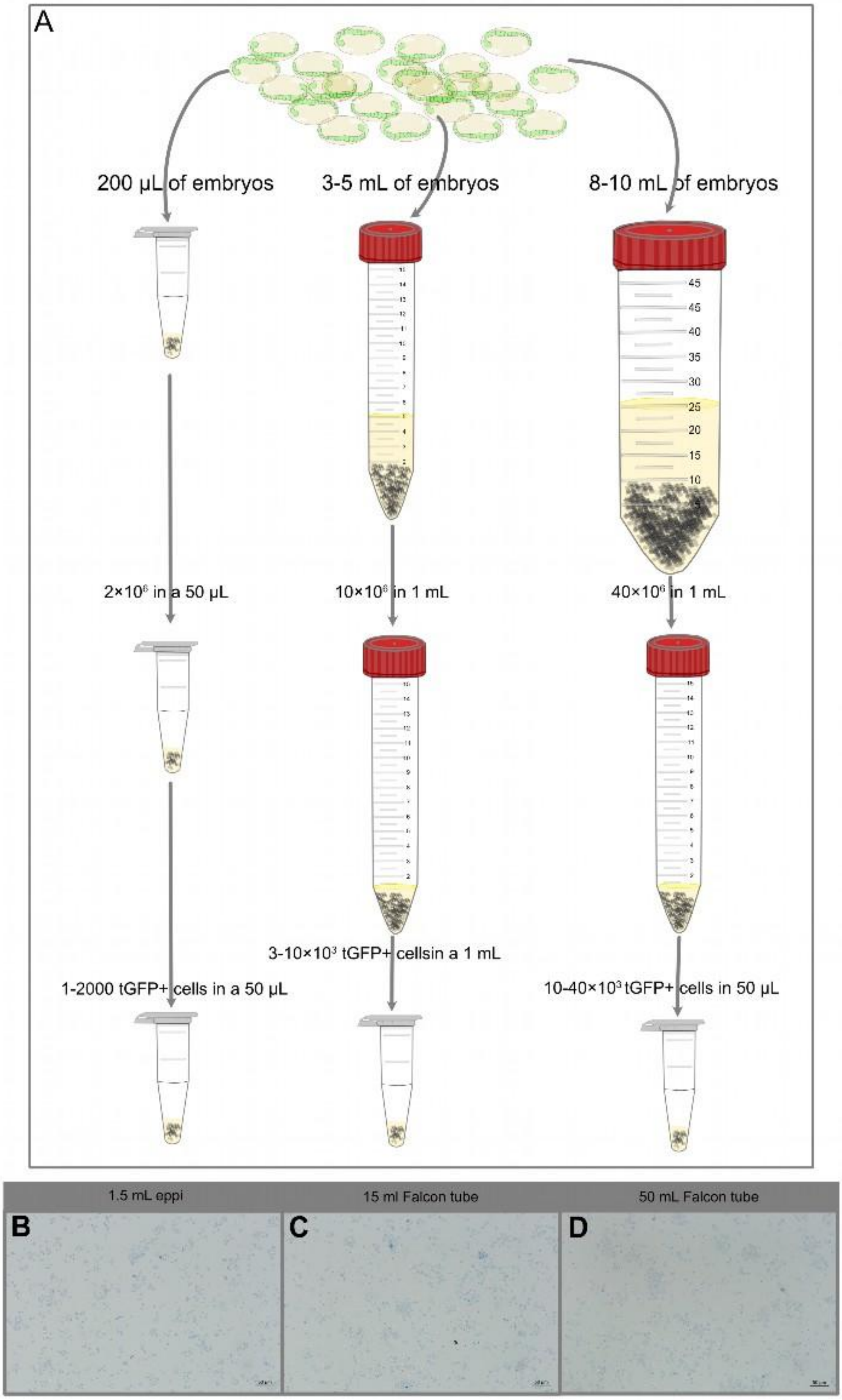
A
200 µL of embryos
3-5 mL of embryos
8-10 mL of embryos
2×10⁶ in a 50 µL
10×10⁶ in 1 mL
40×10⁶ in 1 mL
3-10×10³ tGFP+ cellsin a 1 mL
1-2000 tGFP+ cells in a 50 µL
10-40×10³ tGFP+ cells in 50 µL
1.5 mL eppi
15 ml Falcon tube
50 mL Falcon tube
B
C
D

**Fig. S2. Amount of embryos required for dissociation.** When producing the cell suspensions, we started by using a small number of embryos. However, we escalated our method to reach the number of cells for our experiments. We tested the cell dissociation at different volumes. In **Fig. S2A**, we present three amounts of transgenic embryos and the number of GFP-positive cells recovered after cell sorting. Our methodology required collecting a high amount of GFP -positive cells, but we realized that the yield of these cells after sorting was insufficient. We escalated the method to dissociate a high amount of *Tribolium* embryos. We show that it is possible to obtain cells from a low number of embryos by performing the dissociation in a 1.5 mL microcentrifuge tube (**Fig S2, B**). We can also increase the volume to 5 mL in 15 mL Falcon tubes (**Fig S2, C**) and 30 mL in 50 mL Falcon tubes (**Fig S2, D**). Scale bar 50 μm.

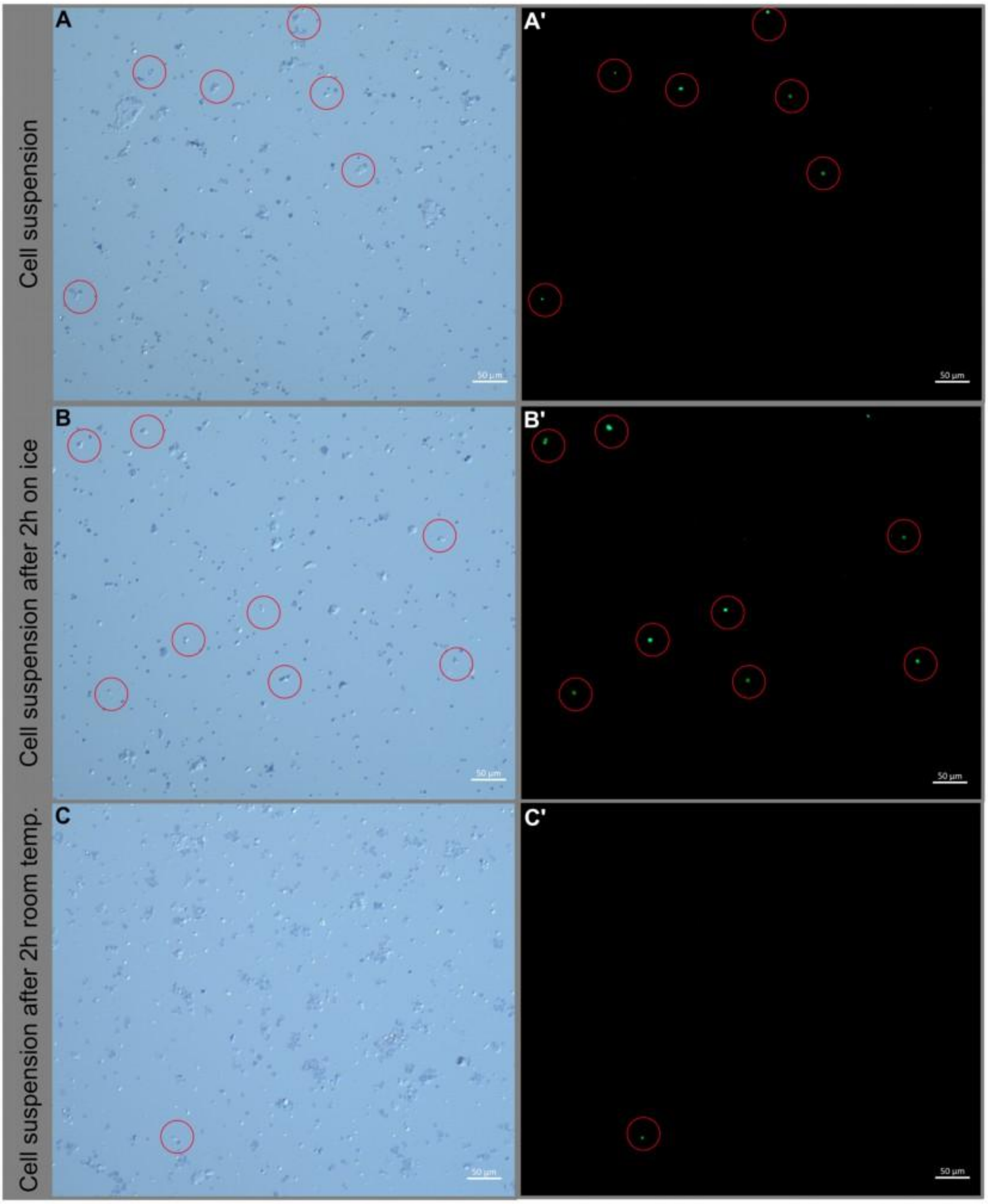
Cell suspension
Cell suspension after 2h on ice
Cell suspension after 2h room temp.
A
A'
B
B'
C
C'
50 µm

**Fig S3.** ***Tribolium*** **cell suspensions maintained at different temperatures.** We tested whether keeping the cell suspensions at different temperatures impacted the survival of the GFP-positive cells. We inspected the cell suspension under the microscope just after cell dissociation (**Fig S3, A** and **Fig S3, A'**) and identified a mix of dead and live cells. We also identified GFP-positive cells (**Fig S3, A'**) marked in the red circles in the same sample. The same cell suspension was divided into two parts, one was placed on ice, and another one was placed at room temperature, both samples were analysed after two hours. The sample kept on ice displayed similar results as the fresh sample. We identified dead and live cells (**Fig S3, B**) and GFP cells (**Fig S3, B'**) marked in the red circles. The sample kept at room temperature showed that cells started to aggregate (**Fig S3, C**), and we saw a decrease in the number of GFP-positive cells (**Fig S3, C'**) shown in the red circles. Scale bar 50 µm.
